# Grid cell disruption in a mouse model of early Alzheimer’s disease reflects reduced integration of self-motion cues and increased influence of environmental geometry

**DOI:** 10.1101/2022.12.15.520595

**Authors:** Johnson Ying, Antonio Reboreda, Motoharu Yoshida, Mark P. Brandon

## Abstract

Grid cell impairments and path integration deficits are sensitive markers of early Alzheimer’s disease (AD). Converging evidence from human and rodent studies suggest that disrupted grid coding underlies path integration deficits in preclinical individuals. However, it still remains unclear if disrupted early AD grid coding reflects increased noise across the network or a specific deficit in path integration, perhaps via an impairment in the integration of self-motion cues. Here, we report in the J20 transgenic amyloid beta mouse model of early AD that grid cells were spatially unstable towards the center of the square arena but not near the borders, had qualitatively different spatial components that aligned parallel to the borders of the environment, and exhibited impaired integration of distance travelled via reduced theta phase precession. Our results suggest that disrupted early AD grid coding reflects reduced integration of self-motion cues but not environmental landmarks, providing further evidence that grid cell impairments underlie specific path integration deficits in early AD.

## Introduction

Grid cells in the medial entorhinal cortex (MEC) fire in multiple spatial locations to form a periodic hexagonal array that spans two-dimensional space (Fyhn et al., 2004; Hafting et al., 2005; Jacobs et al., 2013). This periodic code is implicated in path integration, a cognitive function involving the integration of self-motion cues to maintain one’s sense of location relative to a starting point in space (Gil et al., 2018; McNaughton et al., 2006; Mittelstaedt & Mittelstaedt, 1980; Segen et al., 2022). Both grid cell and path integration impairments are sensitive markers of pathological decline during early AD in human subjects and mouse models of pathology (Bierbrauer et al., 2020; Coughlan et al., 2018; Kunz et al., 2015; Mucke et al., 2000; Segen et al., 2022; Ying et al., 2022).

These early grid impairments may result from disrupted processing of self-motion cues, which constitute necessary inputs that maintain grid representations in healthy animals (Buetfering et al., 2014; Campbell et al., 2018; Chen et al., 2019, 2016; Couey et al., 2013; Hafting et al., 2005; Kraus et al., 2015; Kropff et al., 2015; Miao et al., 2017; Pérez-Escobar et al., 2016; Sargolini et al., 2006; Winter, Clark, et al., 2015; Winter, Mehlman, et al., 2015). Yet, external landmarks also exert significant influences over grid coding, particularly in rectilinear environments where grid representations scale proportionally to manipulations along the borders (Barry et al., 2007). Furthermore, deformed grid hexagonal symmetry in asymmetrical enclosures such as trapezoids demonstrates the degree to which geometric landmarks compete with self-motion cues to generate grid firing (Krupic et al., 2015).

To distinguish if early AD grid cell impairments reflect increased noise across the network or a specific deficit in processing self-motion cues over external landmarks, we analyzed our *in vivo* electrophysiological dataset of MEC neurons recorded in the J20 transgenic amyloid beta (Aβ) mouse model of early AD that expresses a mutant form of human amyloid precursor protein (APP) – referred to here as ‘APP mice’ (Mucke et al., 2000; Ying et al., 2022).

## Results

### Adult APP grid cells are spatially unstable in the center of the environment

We analyzed 4524 MEC neurons from 38 APP transgenic and 30 non-transgenic (nTG) littermates as they foraged for water droplets in a 75 cm square arena (*Summary of MEC recordings,* **Table S1**; *MEC Tetrode locations,* **Figure S1**) (Ying et al., 2022). Mice were recorded between the ages of 3-7 months, timepoints corresponding to the early stages of pathology prior to the expression of widespread Aβ plaques (detailed pathology description in Materials and methods, *Experimental model and subject details*). Mice were categorized into two age groups: young mice between 3-4.5 months of age (APP-y and nTG-y) and aged mice between 4.5-7 months of age (APP-a and nTG-a). Cells with gridness scores (a measure of hexagonal spatial periodicity in the rate map) higher than the 99^th^ percentile of a shuffled distribution (a gridness score above 0.54) were characterized as grid cells (Materials and methods, *Grid cell and head-direction cell selection*).

To quantify spatial stability of grid cells across the environment, we divided the spatial arena into “wall” or “center” regions. The wall length was selected to be 12 cm, corresponding to the body length of a mouse and approximately divides wall and center regions into equal surface areas (**Figure 1A**). Recordings were partitioned into two halves either by time (15 minutes each) or by the animal’s occupancy (Materials and methods, *Spatial stability analysis).* Both methods revealed greater center instability in APP-a grid cells, but wall stability was unchanged across groups (**Figures 1B-D**; n = 36, 64, 30 and 37 cells for nTG-y, nTG-a, APP-y, APP-a). We repeated our analysis in an unbiased manner by incrementally shifting the layer of spatial bins corresponding to both wall and center regions (**Figure S2**). In both time-partitioned and occupancy-partitioned analyses, APP-a center stability was lower across multiple conditions. In contrast, wall stability generally remained unchanged across groups until more center spatial bins were included into the wall region (**Figures S2A and S2B**). Out of all groups, APP-a grid cells also had the most instances where stability was lower in the center versus the borders (**Figure S2C and S2D**). Greater instability towards the environmental center where the sparsity of external landmarks necessitates the use of path integration to maintain a stable grid code suggests that APP-a grid cells do not reliably process self-motion cues.

**Figure 1.**
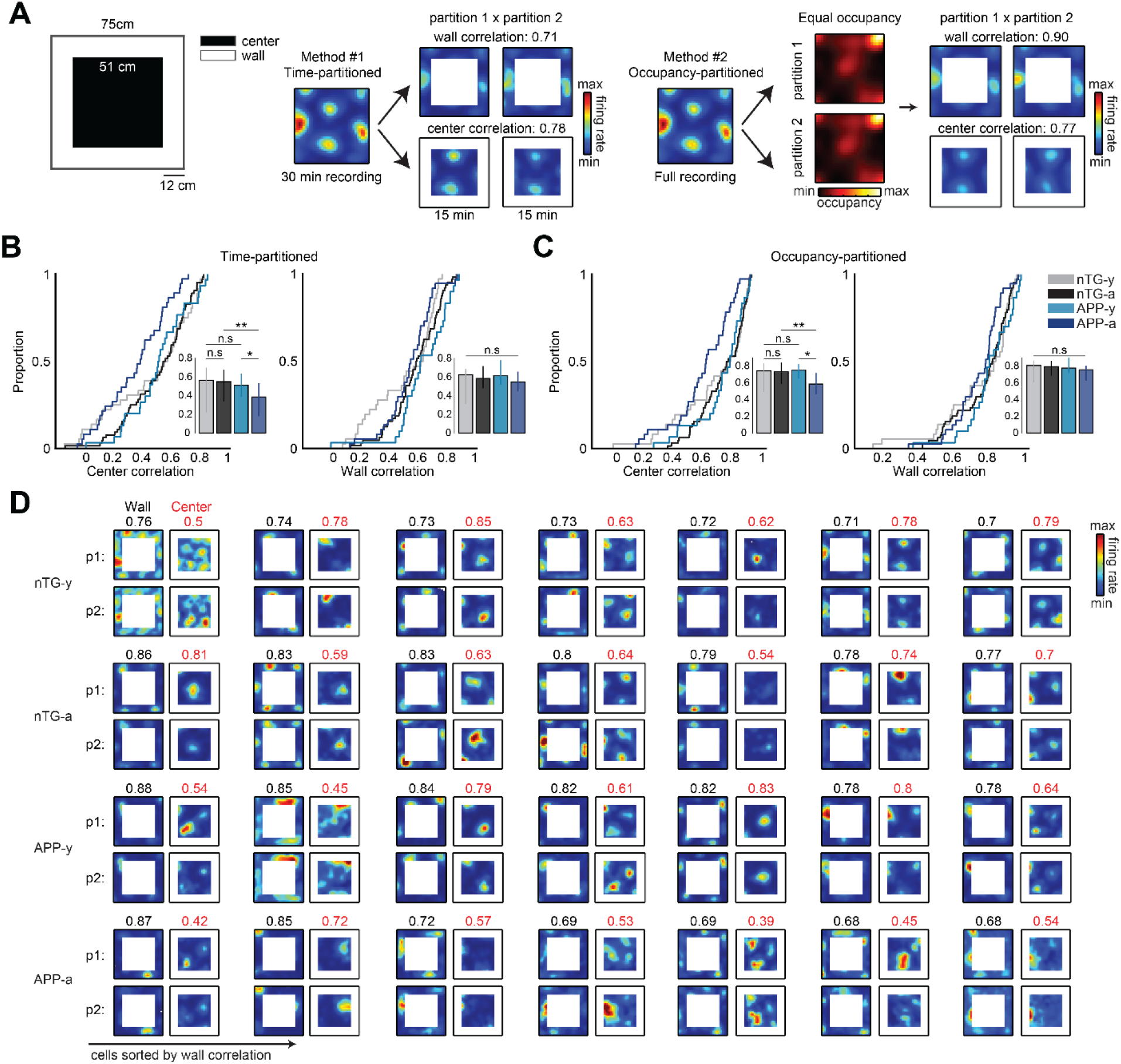
Adult APP grid cells are spatially unstable in the center of the environment. **(A)** The spatial arena was divided into ‘wall’ and ‘center’ regions. The wall length was 12 cm. Spatial stability was calculated via two methods. A time-partitioned analysis split the first 30 minutes of each grid cell recording into two 15 minutes partitions. An occupancy-partitioned analysis split the entire recording into two maps where the occupancies in each spatial bin were the same. Higher correlation scores between wall or center partitions indicate higher spatial stability. **(B)** Center stability is reduced in time-partitioned APP-a grid cell maps, but wall stability is preserved. Bars indicate medians, and error bars show the 25^th^ and 75^th^ percentile, two-way ANOVA (center - age x genotype interaction: F(1, 163) = 7.2, p = 0.008; wall - age x genotype interaction: F(1, 163) = 8.1, p = 0.005), Wilcoxon rank sum post hoc tests with a Bonferroni-Holm correction. **(C)** Center stability is reduced in occupancy-partitioned APP-a grid cell maps. Bars indicate medians, and error bars show the 25^th^ and 75^th^ percentile, two-way ANOVA (center - age x genotype interaction: F(1, 163) = 9.14, p = 0.0029; wall - age x genotype interaction: F(1, 163) = 2.81, p = 0.10), Wilcoxon rank sum post hoc tests with a Bonferroni-Holm correction. **(D)** Time-partitioned rate map examples for seven grid cells with the highest wall stability in each group. Wall and center scores are indicated above each partition (p1 and p2) set in black and red, respectively. Wall correlation scores remained consistently high between groups, but center correlation scores were generally lower in APP-a grid cells. For all panels, n.s = non-significant, *p < 0.05, **p < 0.01.

### Adult APP grid cells exhibit fewer spatial Fourier spectral components

Previously, we reported a disruption in APP-a grid cell hexagonal symmetry (Ying et al., 2022). However, it remains unclear if these disrupted patterns reflect increased random noise in the network or qualitatively different underlying spatial structures. To quantify structural differences of APP-a grid cells, we implemented a two-dimensional Fourier analysis to decompose a cell’s spatial firing rate map into its basic spatial components (Krupic et al., 2012) (Materials and methods, *Generation of Fourier spectrums*). Grid cells typically had three components in the Fourier spectrum which could be visualized as images of spatial axes facing specific orientations offset by 60°, such that the sum of all component images produced the firing rate map (**Figure 2A**). Similar to prior findings, grid cell Fourier components had distinct wavelengths (spacing between axes) where modules scaled by multiples ranging from 1.34 to 1.59 in all groups (**Figure S3**) (Barry et al., 2007; Krupic et al., 2012; H. Stensola et al., 2012). To discount the inherent biases of grid cell selection criteria, we also applied the Fourier analysis to all recorded MEC cells. Most non-grid MEC cells typically had two Fourier components and adopted a quadrant-like arrangement of spatial axes with angular offsets being multiples of 90° (**Figure 2A**; n = 768, 997, 720 and 1318 cells for nTG-y, nTG-a, APP-y, APP-a). We computed the polar autocorrelations of all recorded MEC cells by circularly shifting each cell’s polar representation by 360° and found a noticeable decrease in 60° modulation but an increase in 90° modulation for APP-a mice (**Figure 2B**). These results suggest that APP-a spatial axes at the population level were less hexagonal and more quadrant-like.

**Figure 2.**
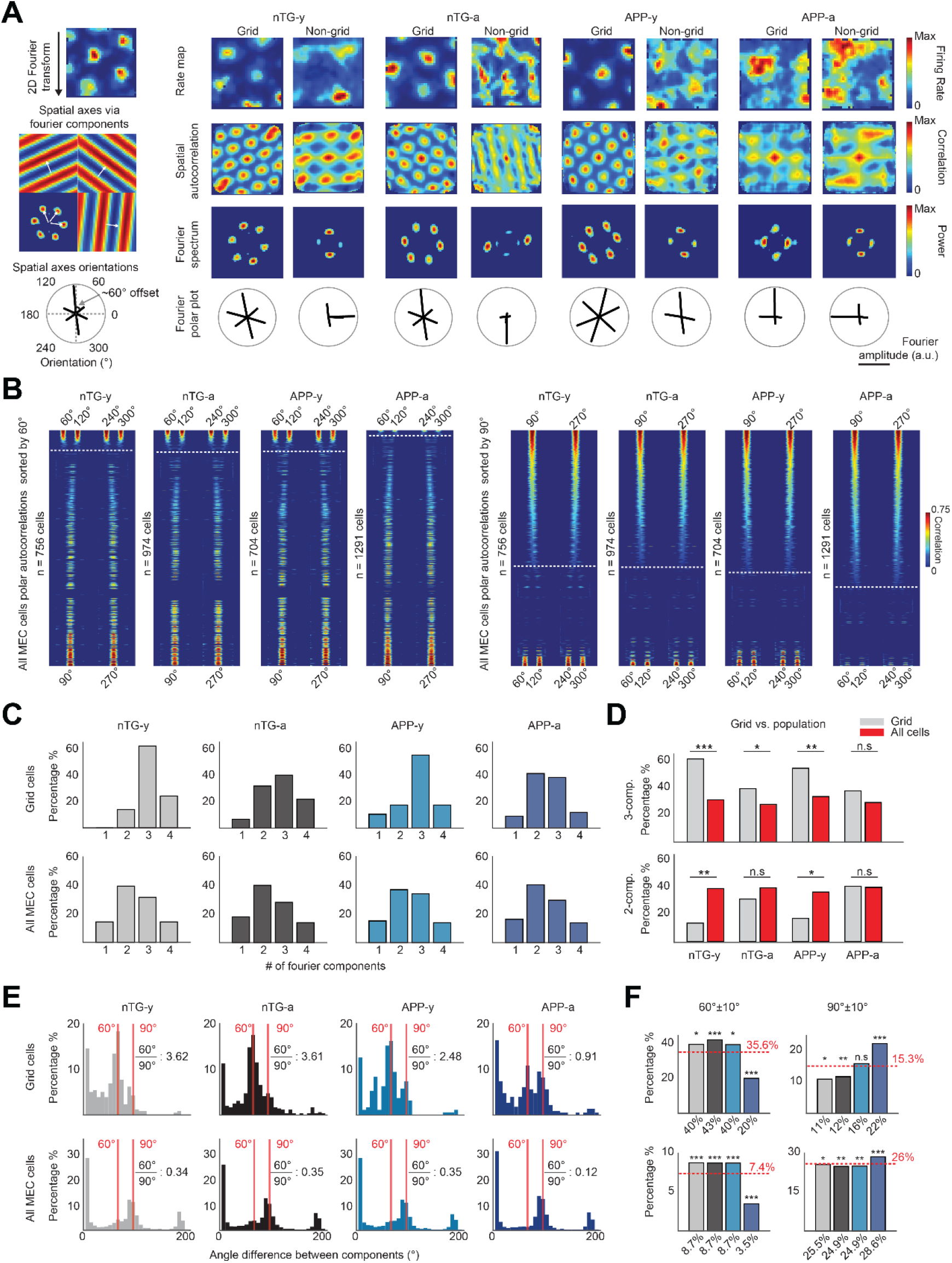
Adult APP MEC neurons more frequently adopt quadrant-like spatial alignment. **(A)** A 2D Fourier transform decomposes the rate map into a small number of Fourier components. Each grid cell component can be visualized as an image of grid axes typically oriented at multiples of 60°. The orientations of all Fourier components could be visualized as a polar plot. Single grid and non-grid spatially periodic cell examples are shown for all groups. **(B)** Polar autocorrelations for all MEC cells sorted by the strength of hexagonal modulation at 60° multiples or by the strength of quadrant-like modulation at 90° multiples. Dashed white lines indicate where hexagonal modulation or quadrant-like modulation ends. **(C)** Percentage distribution of cells that had one-to-four Fourier components for grid cells and all MEC cells. **(D)** The percentages of grid cells that had either three (one-tailed t-test for proportions) or two components (two-tailed t-test for proportions) versus all MEC cells. **(E)** Histograms show the distribution of angle difference between neighboring components of grid and all MEC cells. Solid red lines mark specific orientations. **(F)** Bar graphs show the percentages of angular differences at 60°±10° and 90°±10° compared to an expected percentage calculated as the average of observed proportions across all groups (dotted red line). Binomial test. For all panels, n.s = non-significant, *p<0.05, **p <0.01, ***p< 0.001.

Further quantification of Fourier component count revealed that APP-a mice had less 3-component grid cells. Grid cells in nTG-y, nTG-a and APP-y mice generally had three significant Fourier components, while grid cells in APP-a mice had a similar number of cells with two or three components (**Figure 2C**). Most non-grid MEC cells had two significant Fourier components (**Figure 2C**). There was a higher percentage of 3-component grid cells than 3-component MEC cells in all groups except for APP-a mice (**Figure 2D**; nTG-y: p < 0.001, nTG-a: p = 0.024, APP-y: p = 0.01, APP-a: p = 0.14). On the other hand, there was a higher percentage of 2-component MEC cells than 2-component grid cells in nTG-y and APP-y mice, but not in nTG-a and APP-a mice (**Figure 2D**; nTG-y: p = 0.005, nTG-a: p = 0.20, APP-y: p = 0.03, APP-a: p = 0.93). These results suggest that only APP-a mice had both a larger percentage of 2-component grid cells and a smaller percentage of 3-component grid cells.

To determine the type of spatial alignment adopted by these 2- and 3-component cells, we computed the angular difference of neighboring Fourier components. Angular offsets in grid cell neighboring components were mostly 60° (and occasionally 90°) in nTG-y, nTG-a, and APP-y mice, but we observed similar percentages between 60° and 90° offsets in APP-a mice (**Figures 2E and 2F**; 60°±10°: nTG-y: 40%, nTG-a: 43%, APP-y: 40%, APP-a: 20%; 90°±10°: nTG-y: 11%, nTG-a: 12%, APP-y: 16%, APP-a: 22%). Furthermore, the ratio of 60° to 90° was substantially reduced in APP-a grid cells (**Figure 2E**; (60°±10°)/(90°±10°): nTG-y: 3.62 nTG-a: 3.61 APP-y: 2.48 APP-a: 0.91). Compared to chance level (the average of observed percentages across all groups), only APP-a mice had significantly less instances of 60° but more of 90° (**Figures 2E and 2F**; APP-a vs. 60° chance: p = 1.13 x 10^-7^; APP-a vs. 90° chance: p = 9.98 x 10^-4^). These results suggest that APP-a grid cell axes deviate from a 3-component hexagonal alignment and more strongly adopt a 2-component quadrant-like alignment. Angular differences in the neighboring components of all MEC cells were predominantly multiples of 90° in all groups (**Figure 2E**). Relative to other groups, APP-a MEC cells had a ~60% decrease in 60° angular offsets and a ~15% increase in 90° angular offsets (**Figure 2F**; 60°±10°: nTG-y: 8.7%, nTG-a: 8.7%, APP-y: 8.7%, APP-a: 3.5%; 90°±10°: nTG-y: 25.5%, nTG-a: 24.9%, APP-y: 24.9%, APP-a: 28.6%).

### Adult APP grid cell spatial axes align parallel to environmental borders

Our finding that the spatial stability of grid cells was reduced towards the arena’s center but not near the borders suggests that APP-a grid cells remained anchored to the environment despite a potential impairment of self-motion cue integration. To quantify this anchoring, we analyzed how hexagonal and quadrant-like spatial codes aligned to the environment’s geometry. Most 2-component and 3-component cells adopted one of three alignment profiles (**Figure 3A**). Consistent with previous reports, many 3-component cells were either 30° or 60° hexagonally offset from the east wall (**Figure 3A**) (Krupic et al., 2012; H. Stensola et al., 2012; T. Stensola et al., 2015). On the other hand, most 2-component cells were aligned parallel to the borders at multiples of 90° (**Figure 3A**). APP-a mice had ~8% more MEC cells that adopted this quadrant-like alignment profile and 5-6% less cells that adopted hexagonal alignment profiles (**Figure 3A**; quadrant-like cells: p = 5.47 x 10^-10^, hexagonal cells: p = 4 x 10^-8^).

**Figure 3.**
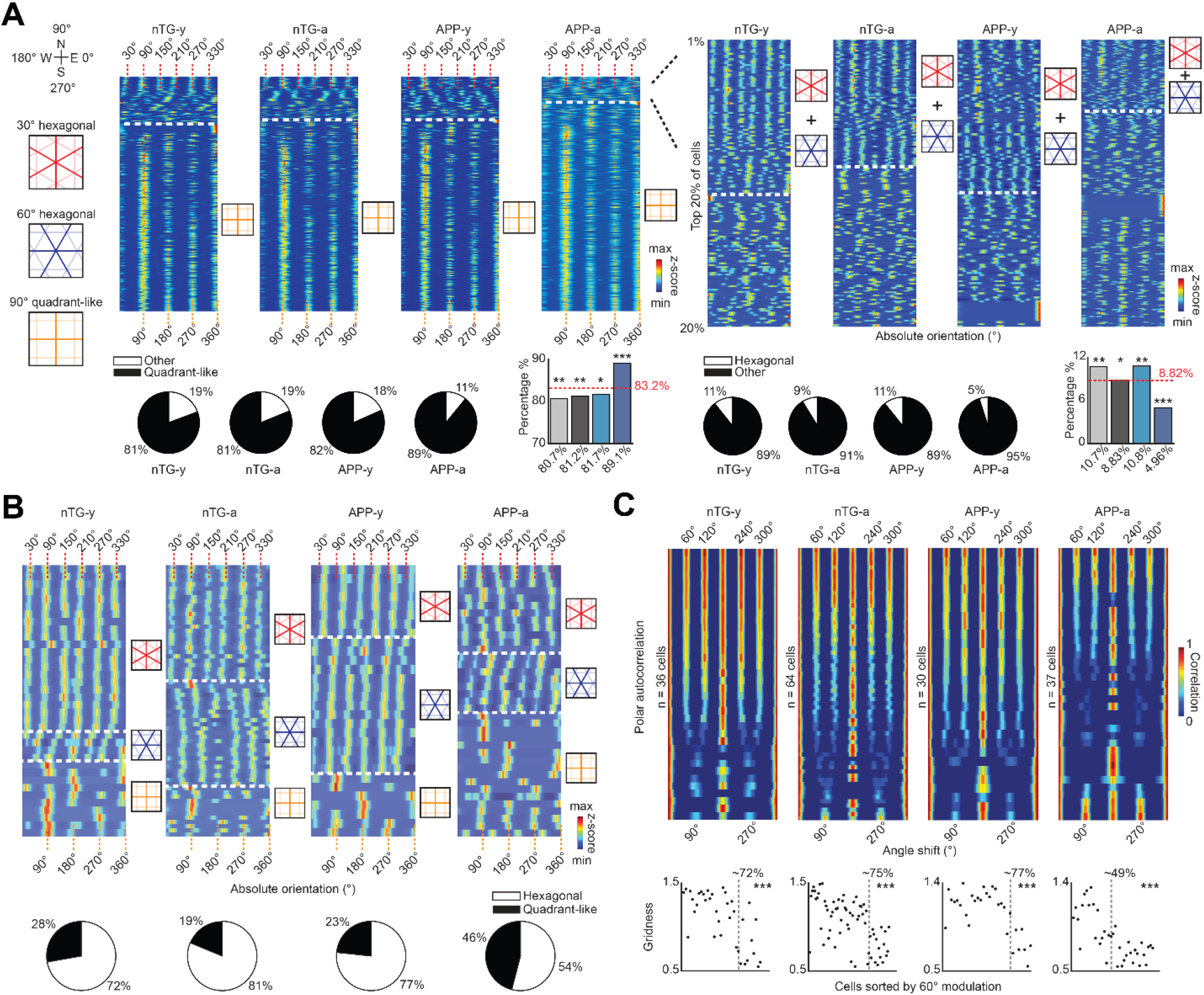
Adult APP grid cells align parallel to the borders. **(A)** (Left-top) Absolute orientation of all cells relative to the recording environment. Dashed white lines indicate where quadrant-like alignment starts. (Left-bottom) Percentage distributions of quadrant-like cells between groups, and compared to the expected chance of recording them (dotted red line). Binomial test. (Right-top) Zoomed-in view of the absolute orientations of the top 20% of cells in the left panel. Dashed white lines indicate where 30° or 60° hexagonal alignment ends. (Right-bottom) Percentage distributions of hexagonal cells between groups, and compared to the expected chance of recording them (dotted red line). Binomial test. **(B)** Top: Absolute orientation of all grid cells relative to the recording environment. Grid cells could be grouped into 30° or 60° hexagonal alignment, as well as 90° quadrant-like alignment (boundaries separated by black dotted lines). To account for differences in Fourier power between cells, colors indicate the z-score of each polar representation. Bottom: Percentage distributions of hexagonal or quadrant-like grid cells between groups. **(C)** Polar autocorrelation for grid cells in all groups. Grid cells are sorted by the strength of hexagonal modulation at 60° multiples. Colors indicate correlation at each degree shift along the x-axis. Bottom: Gridness scores of grid cells sorted by their degree of hexagonal or quadrant-like modulation (90° intervals) along the x-axis. The boundary between the two categories is indicated by the gray dotted lines, and the percentage of strongly hexagonally modulated grid cells are indicated for each group. Wilcoxon rank sum tests. For all panels, n.s = non-significant, *p < 0.05, **p < 0.01, ***p < 0.001.

Prior to visualizing the spatial alignment of grid cells, we considered the possibility that many 1- or 2-component grid cells may have been falsely classified 3-component grid cells as a result of Fourier components not being properly identified in the Fourier spectrum due to relaxed image detection parameters. To control for this possibility, we increased image detection thresholds to obtain an additional 29% (4/14 cells), 54% (14/26 cells), 46% (6/13 cells) and 33% (9/27 cells) 3-component grid cells from 1- and 2-component grid cells in nTG-y, nTG-a, APP-y, and APP-a mice, respectively (Materials and methods; **Figure S4**). From this corrected data set, we observed that all groups had 30° and 60° 3-component grid cells, or 90° 2-component grid cells (**Figure 3B**). Specifically, we found 28% (10/36 cells), 19% (12/64 cells), 23% (7/30 cells), and 46% (17/37 cells) 1- and 2-component grid cells aligned parallel to the borders in nTG-y, nTG-a, APP-y, and APP-a mice, respectively. APP-a mice had a greater percentage of 1- and 2-component grid cells by ~20-25% than other groups (**Figure 3B**; **Table S2**; p = 0.01). Polar autocorrelations revealed that ~75% of grid cells were modulated at 60° intervals in nTG-y, nTG-a and APP-y mice, whereas ~25% of cells were modulated at 90° intervals (**Figures 3C and S5)**. In contrast, APP-a grid cells were roughly equally split between 60° and 90° modulation (**Figure 3C**). Notably, 3-component grid cells had higher gridness scores than their 1- and 2-component counterparts in all groups (**Figure 3C**). These results demonstrate that our previously reported finding of disrupted grid cell hexagonal symmetry in APP-a mice is not due to increased random noise, but rather caused by a higher concentration of firing aligned parallel to the borders (Ying et al., 2022). We confirmed that our results are not artifacts caused by the parameters of our correction threshold to retrieve 3-component grid cells or the square dimensionality of rate map images (**Table S3 and Figure S6**).

### Adult APP grid cells exhibit reduced theta modulation and positional coding via theta phase precession

Given the importance of theta rhythmicity for grid cell spatial coding (Brandon et al., 2011; Koenig et al., 2011) and the proposed link between entorhinal or hippocampal theta rhythms and various forms of self-motion including running speed, positive acceleration and vestibular inputs (Hafting et al., 2008; Jacob et al., 2014; Jeewajee et al., 2008; Kropff et al., 2021; Maurer et al., 2005; O’Keefe & Recce, 1993; Ravassard et al., 2013; Skaggs et al., 1996; Terrazas et al., 2005; Winter, Clark, et al., 2015), we examined whether intrinsic theta rhythmicity of spiking was reduced in APP-a grid cells. Visualization of the spike–time autocorrelations of grid cells revealed weaker overall theta modulation than other groups (**Figure 4A**). Visualization of the power spectrums of the spike-time autocorrelations yielded a similar conclusion. APP-a grid cells showed weaker theta modulation around 8-10 Hz compared to other groups (**Figure 4B**). Next, we quantified the percentages of theta-modulated grid cells from the power spectrums of the spike-time autocorrelations (Materials and methods, *Theta modulation analyses*). There were 21-31% less theta-modulated APP-a grid cells compared to other groups (p = 0.009) (**Figure 4C**; nTG-y: 69%, nTG-a: 77%, APP-y: 67%, APP-a: 46%). We repeated our quantifications in an unbiased manner for a wide range of temporal bin sizes between 1-10 ms and observed a consistent ~21-31% reduction in theta-modulated APP-a grid cells than other groups (**Figure 4D**).

**Figure 4.**
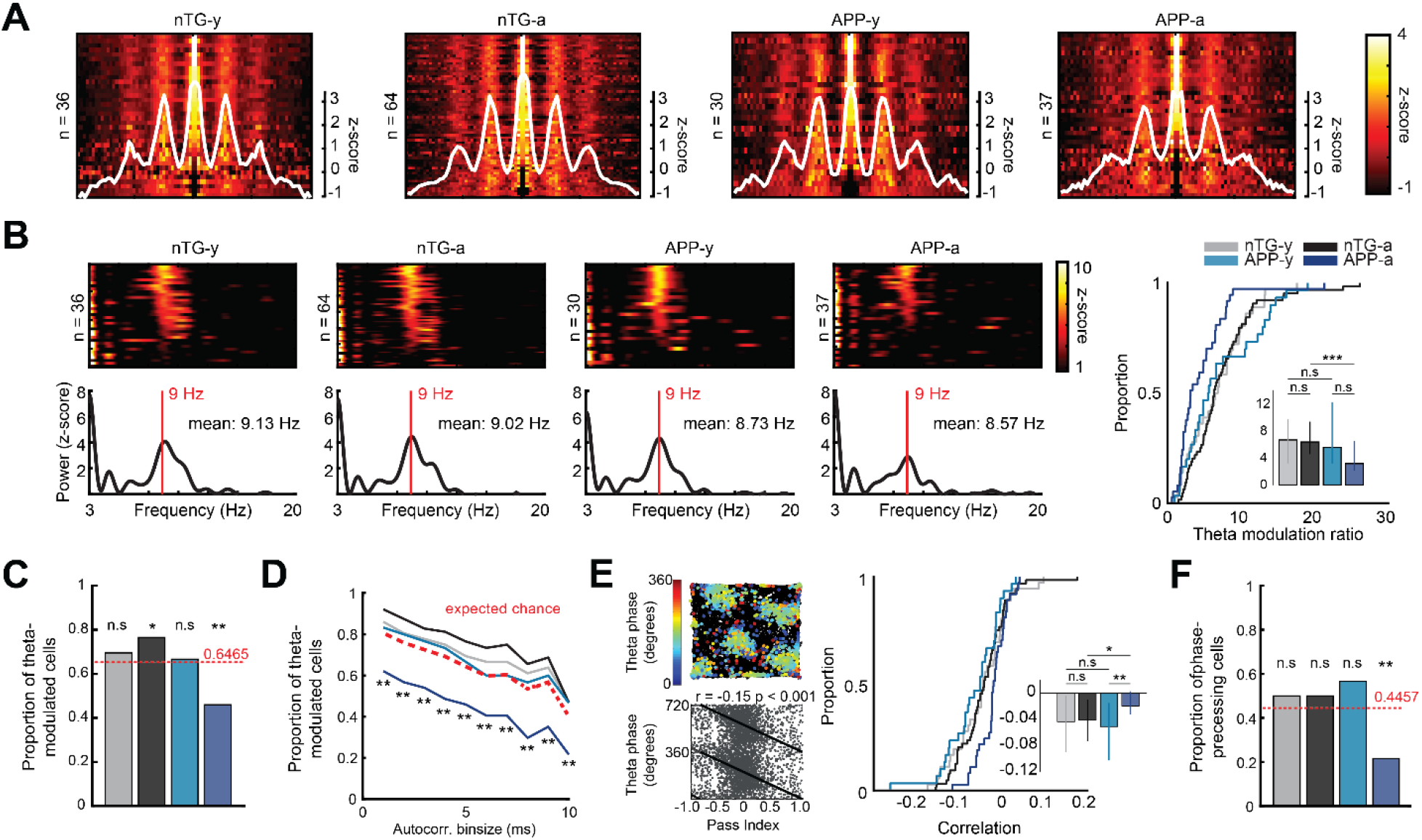
Adult APP grid cells have reduced theta modulation and theta phase precession. **(A)** Color-coded raster plots of spike-time autocorrelations for grid cells across groups. White lines are the average autocorrelation curves. **(B)** Left: Color-coded raster plots show the power spectrums of spike-time autocorrelations for grid cells across groups. Line graphs show the mean intrinsic frequency curve and red lines mark where a 9 Hz intrinsic frequency. Right: Theta modulation ratio is reduced in APP-a grid cells. Bars indicate medians, and error bars show the 25^th^ and 75^th^ percentile, two-way ANOVA (age x genotype interaction: F(1, 163) = 5.33, p = 0.022), Wilcoxon rank sum post hoc tests with a Bonferroni-Holm correction. **(C)** Proportions of significantly theta-modulated grid cells compared to the expected chance of recording them (dotted red line). Binomial test. **(D)** Repetition of **(C)** when varying the bin size of the spike-time autocorrelation between 1 to 10 ms. In all conditions, there was a consistent decrease in significantly theta-modulated grid cells in APP-a mice. **(E)** Left: Open field phase precession for an example grid cell. Trajectory of mouse in black with overlaid color-coded circles indicate the location and theta phase of spiking. Pass index values of −1 and +1 represent the entry and exit of a firing field, respectively. Right: The strength of correlation between the spiking phase and distance travelled across a firing field is reduced in APP-a mice. Data includes quadrant-like and hexagonal grid cells. Bars indicate medians, and error bars show the 25^th^ and 75^th^ percentile, two-way ANOVA (age x genotype interaction: F(1, 163) = 5.34, p = 0.02), Wilcoxon rank sum post hoc tests with a Bonferroni-Holm correction. **(F)** Proportion of significantly phase-precessing grid cells compared to the expected chance of recording them (dotted red line). Binomial test. For all panels, n.s = non-significant, *p < 0.05, **p < 0.01.

A potential consequence of non-theta-modulated grid cells is their inability to encode distance travelled via a temporal code known as theta phase precession. As the animal travels across a firing field, a theta-modulated grid cell spikes at progressively earlier phases of theta oscillations in the local field potential (Hafting et al., 2008; O’Keefe & Recce, 1993). Oscillatory interference and continuous attractor network models suggest that theta phase precession properties could allow grid cells to integrate spatial displacement on the basis of self-motion while planning future paths as part of a path integration system (Burgess, 2008; Burgess et al., 2007; Navratilova et al., 2012). To examine theta phase precession of grid cells across groups, we computed the strength of correlation between distance travelled across a field and spike theta phase, and observed an impairment in APP-a mice (**Figure 4E**). If theta modulation is a requirement for theta phase precession, then APP-a grid cells would logically exhibit less phase precession than other groups. Indeed, further quantification revealed that there were 28-35% less phase-precessing APP-a grid cells compared to other groups (p = 0.0022) (**Figure 4F**; nTG-y: 50%, nTG-a: 50%, APP-y: 57%, APP-a: 22%). Together, these results suggest that APP-a grid cells do not reliably integrate self-motion via precisely-timed theta-related mechanisms.

## Discussion

Our results suggest that early grid cell disruption in the J20 mouse model of amyloidopathy reflects reduced integration of self-motion cues and increased influence of environmental geometry. Reduced spatial stability towards the center but not near the borders suggests that grid cell impairments predominantly arise from disrupted processing of self-motion. In parallel, reduced theta modulation and theta phase precession suggests that APP-a grid cells could not properly integrate self-motion cues or accurately encode positional information within theta cycles in the local field potential. In contrast, APP-a grid cells appeared to be more strongly influenced by environmental geometry. A Fourier spectral analysis revealed that disrupted grid spatial periodicity in APP-a mice was not as random as previously assumed (Ying et al., 2022), but directly explained by the degree of spiking aligned parallel to the borders. Grid cell patterns are hypothesized to arise from intrinsic network activity and then anchor to the outside world via external landmarks (Buetfering et al., 2014; Couey et al., 2013; Gardner et al., 2019; T. Stensola & Moser, 2016; T. Stensola et al., 2015; Trettel et al., 2019). The strongest grid anchor appears to be the environmental geometry itself, as different enclosure shapes affect grid cell hexagonal symmetry (Krupic et al., 2015; T. Stensola & Moser, 2016; T. Stensola et al., 2015). Future experiments should investigate if hexagonal and quadrant-like spatial alignment profiles also persist in other types of environmental geometries that vary in polarisation and symmetry. Given that grid hexagonal symmetry is disrupted in trapezoidal geometries (Krupic et al., 2015), one might intuit that MEC spatial patterns will not be as definable as those observed in a square environment. Therefore, we emphasize that the quadrant-like grid firing reported here are limited to square recording environments until there is further evidence in non-square geometries.

What are the implications for reduced theta modulation and phase precession in adult APP mice? The oscillatory interference model posits that one mechanism by which grid cells integrate self-motion is interference of theta oscillations between upstream velocity-controlled oscillators (VCOs) (Burgess, 2008; Burgess et al., 2007; Hasselmo & Brandon, 2008). Different VCO firing frequencies vary according to the animal’s movement speed along angular offsets of 60°. When multiple theta-modulated VCOs with preferred directions evenly spaced around 360° oscillate in phase, the thresholded sum of their directional interference patterns in a band-like manner produces grid hexagonal periodicity and theta phase precession. Alternatively, phase precession is proposed to be a ‘look-ahead’ mechanism to plan future routes and has been modeled in a continuous attractor network (Navratilova et al., 2012). Both oscillatory and continuous attractor models could form the foundations of a path integration system that allows for continuous tracking of position along directions offset by 60° and computation of translational vectors toward goal locations. For instance, neural network implementations of grid cells can be successfully trained to produce vector-based navigation by utilizing phase precession properties (Bush et al., 2015). Specifically, the phase difference relationships of different grid cells encoding current and goal locations can accurately produce goal-directed translational vectors in large-scale two-dimensional spaces (Bush et al., 2015). The lack of phase precession in many APP-a grid cells could impair their ability to integrate self-motion and plan future paths in the environment’s center, thus causing the grid cell network to adopt quadrant-like alignment that may integrate spatial displacement through other means such as visual cues and contact with environmental borders. The sequential organization of grid cell spikes within continuous theta cycles might also constitute a temporal readout of movement direction within short time windows (Zutshi et al., 2017). Theta phase precession suggests that past grid fields fire at earlier phases in a theta cycle while recent grid fields fire at later phases. Each theta cycle therefore describes past, current and future locations. This continuous phase code could underlie the animal’s ability to infer position relative to a starting location when path integrating across long behavioral time scales. Phase precession has also been reported in other species including bats and humans, suggesting that phase coding serves a broad role in linking sequential locations and events (Eliav et al., 2018; Qasim et al., 2021).

Many real-world and virtual human path integration behavioral paradigms eliminate or control against the use of environmental borders (Allen et al., 2004; Bierbrauer et al., 2020; Howett et al., 2019; Mahmood et al., 2009; Mokrisova et al., 2016; Stangl et al., 2020). Partial or complete removal of external landmarks could force preclinical individuals to rely on self-motion information which they have difficulty integrating, thus leading to unstable and compromised fMRI grid-like signals (Bierbrauer et al., 2020; Kunz et al., 2015; Segen et al., 2022). Hexagonal and quadrant-like alignment may suggest general principles about how one encodes position via self-motion. Hexagonal structure is superior to quadrant-like structure in terms of angular resolution and sampling frequency between vertices (Mersereau, 1979). In the context of path integration, 60° grid axes allow for more frequent updating of heading direction, as well as spatial displacement or movement speed between intersection points. These advantages may explain why grid cells in general adopt hexagonal symmetry, or why grid-like representations are modulated at 60° and not at 90° intervals (Doeller et al., 2010).

Tauopathy and amyloidopathy are pathological hallmarks of AD (Berron et al., 2021; Braak & Braak, 1991). While tauopathy is generally regarded as the main driver of entorhinal dysfunction in the general population, different tau and Aβ mouse models independently show a shared disruption of grid cell hexagonal symmetry during early and late pathogenesis (Archetti et al., 2019; Braak & Braak, 1991; Fu et al., 2017; Johnson et al., 2016; Jun et al., 2020; Ossenkoppele et al., 2016; Ridler et al., 2020; Ying et al., 2022; Young et al., 2014). In a two-dimensional continuous attractor neural network model of grid cell activity, simulated AD synaptic damage resulting from the propagation of neurofibrillary tau tangles disrupted grid cell hexagonal symmetry (Zhi & Cox, 2021). Similar to APP-a mice, simulated healthy grid cells had three significant Fourier components offset by 60°. In contrast, simulated damaged grid cells had two, one, or no components depending on the magnitude of synaptic impairment. The similarities between these model simulations of tau propagation and our experimental results in an amyloid mouse model suggest that despite the different molecular pathways of tauopathy and amyloidopathy, the loss of grid hexagonal symmetry across multiple AD mouse models might initially occur through a similar process where the grid map detaches from the individual’s self-motion while staying anchored to the external world. More broadly, our findings support existing theories which suggest that disrupted processing of self-motion by grid cells underlies path integration impairments reported during early AD (Bierbrauer et al., 2020; Howett et al., 2019; Kunz et al., 2015; Segen et al., 2022; Ying et al., 2022).

## Acknowledgements

We graciously thank S. Kim, Z. Ante, K. Harandian, Q. He, A. Ismailova, D. Patel, A. Zhen, and A. Milette-Gagnon for their assistance in experiments. We also thank A.T. Keinath, H.C. Yong, J.Q. Lee, J.C. Robinson and M. Oulé for providing helpful comments on prior versions of the manuscript, as well as all members of the Brandon laboratory for helpful discussions. This work was funded by CIHR Project Grants #367017 and #377074, an NSERC Discovery Grant #74105, a Scottish Rite Charitable Foundation Grant, a Canada Fund for Innovation Grant, and a Canada Research Chairs award to M.P.B. This work was also supported by the German Research Foundation (DFG) project YO177/7-1 and YO177/8-1 to M.Y. J.Y is supported by a Doctoral Training Grant from the Fonds de recherche du Québec, and previously by a Master’s Training Grant from the Fonds de recherche du Québec and a CIHR Master’s Training Fellowship.

## Author contributions

J.Y., A.R., M.Y., and M.P.B. conceived the project and wrote the manuscript. J.Y conducted all experiments. J.Y. and A.R. conducted analysis of data.

## Supplemental Figures

**Figure S1.**
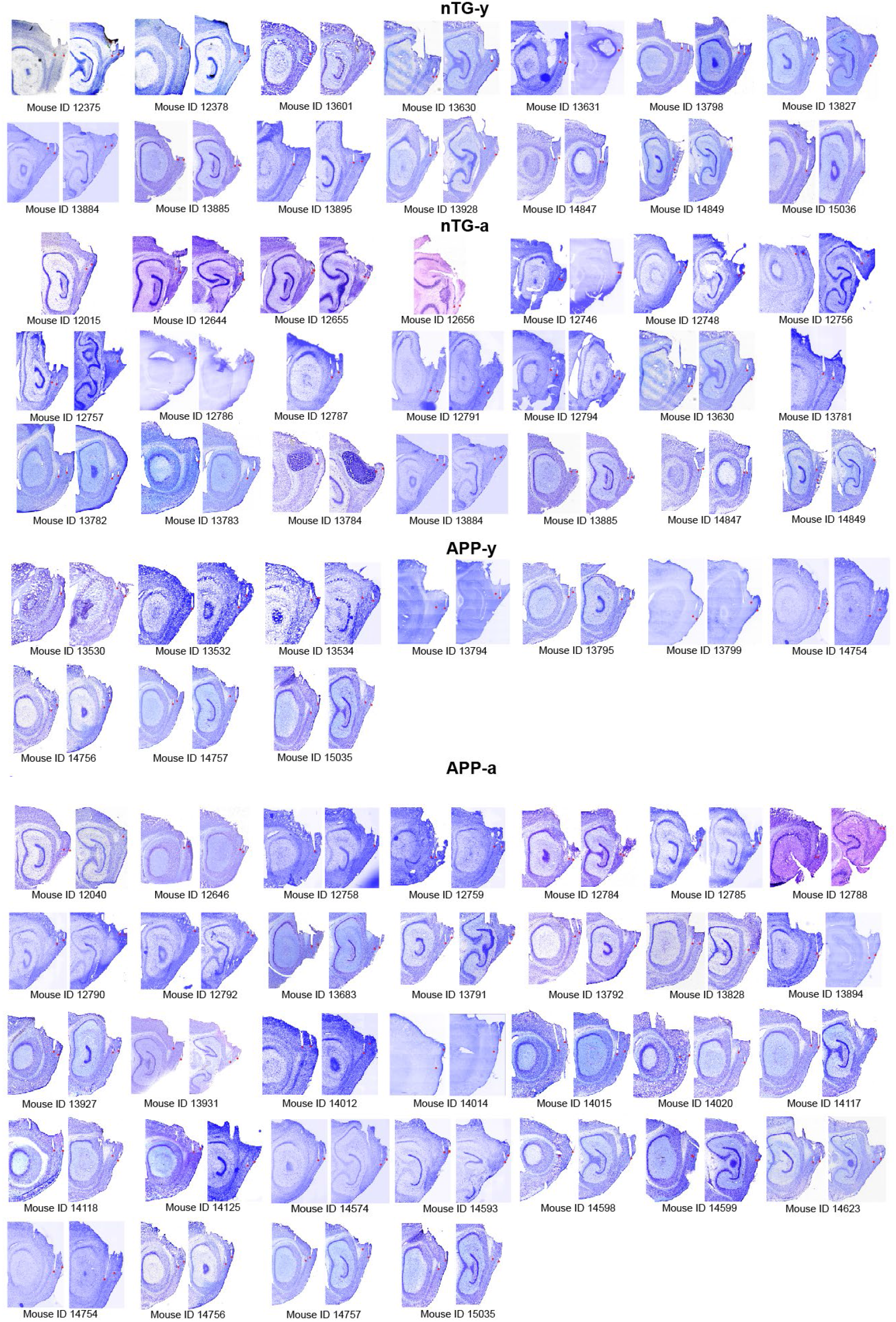
Tetrode track locations. Track locations of each animal ordered by genotype and age. The tips of tracks are highlighted with a red dot.

**Figure S2.**
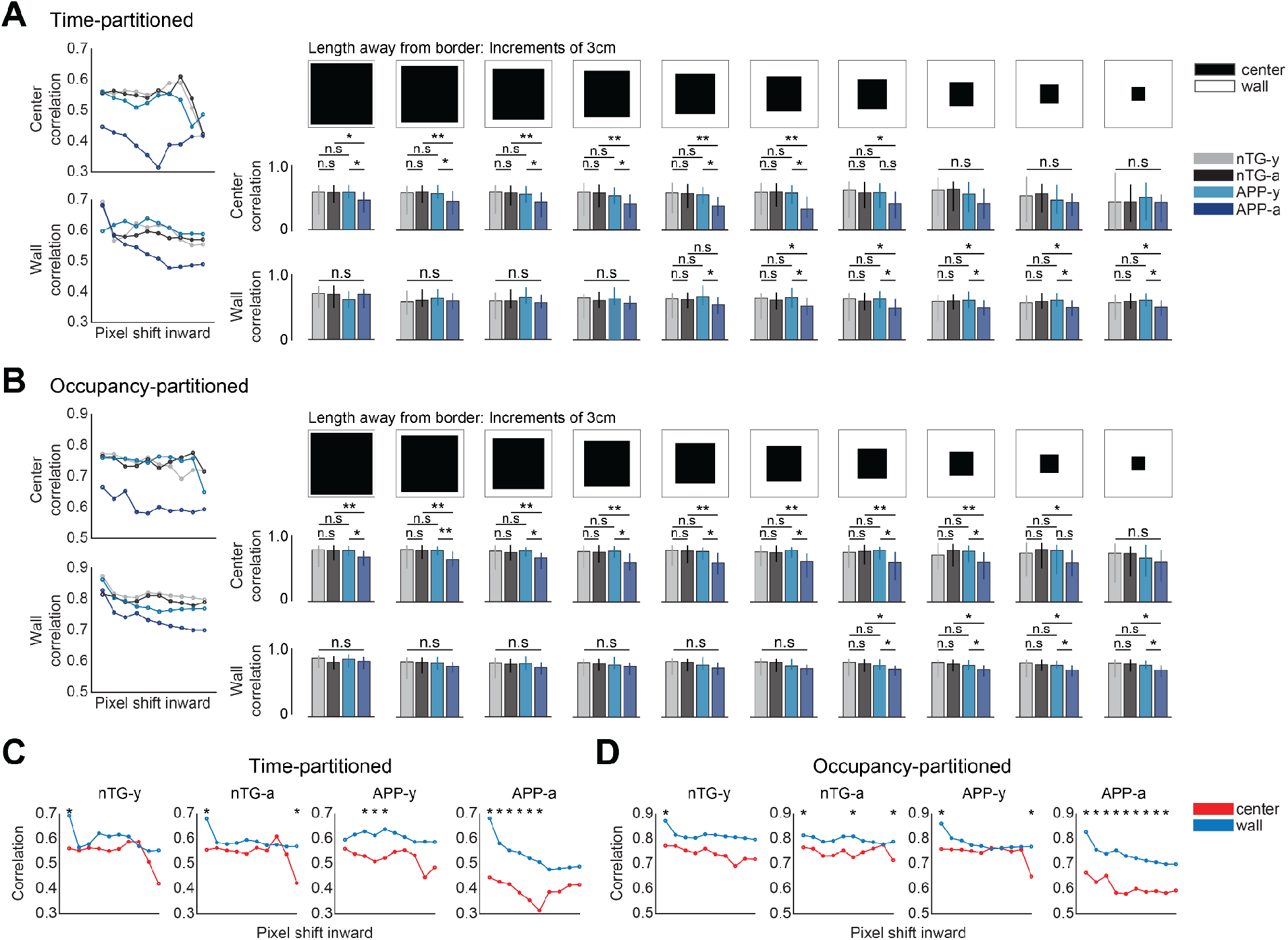
Unbiased spatial stability analyses for grid cells. **(A)** Center or wall Pearson correlations of time-partitioned maps as a relationship of pixel layer shift towards the absolute center. Left: Dots indicate medians. Right: Bars indicate medians, and error bars show the 25^th^ and 75^th^ percentile, two-way ANOVA, Wilcoxon rank sum post hoc tests with a Bonferroni-Holm correction. **(B)** Same as A, but for occupancy-partitioned maps. **(C)** Center versus wall stability of time-partitioned maps as a relationship of pixel shift. Wilcoxon rank sum tests. Stars indicate significant comparisons where p is at least less than 0.05. **(D)** Same as C, but for occupancy-partitioned maps. For panels A-B, n.s. = non-significant, *p < 0.05, **p < 0.01.

**Figure S3.**
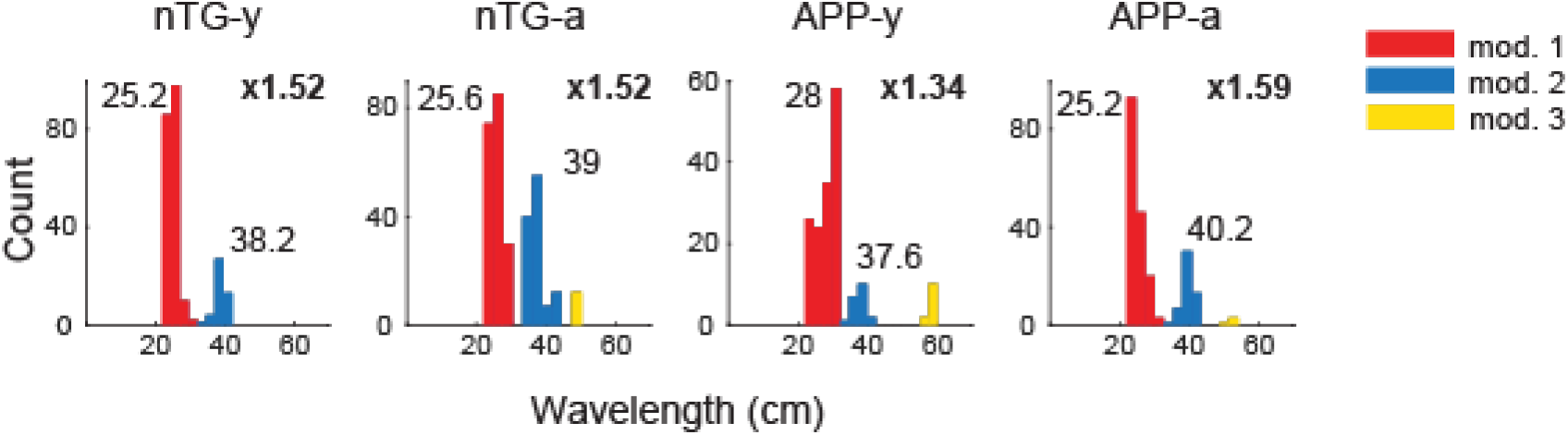
Fourier component wavelengths. Distribution of Fourier component wavelengths for grid cells Colors indicate different wavelength modules. The mean wavelength values for the red and blue modules are shown above, as well as the ratio between modules.

**Figure S4.**
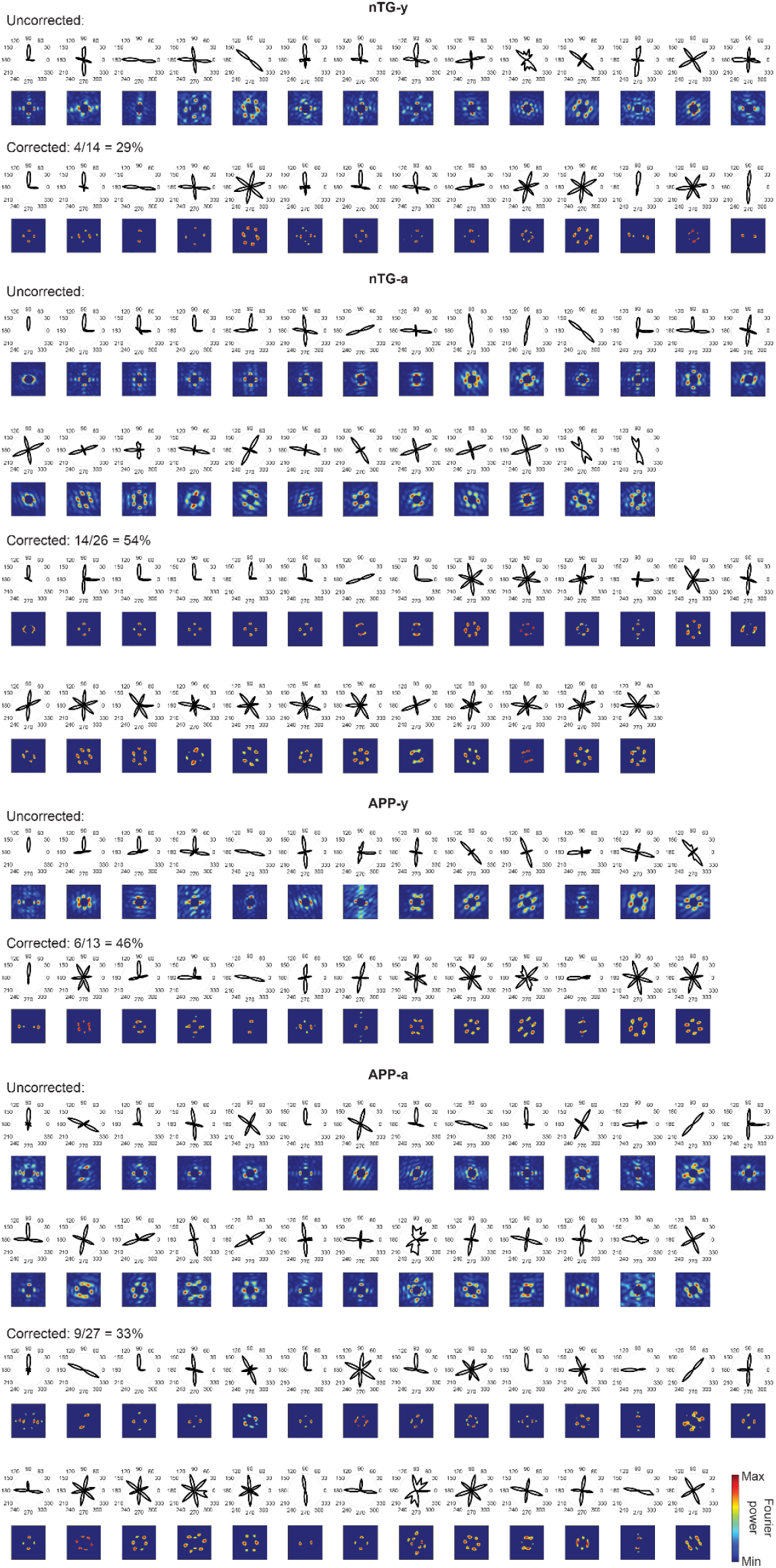
Retrieval of 3-component hexagonal grid cells via correction. All 1-component or 2-component grid cells in the uncorrected group are shown by its polar representation and the raw Fourier spectrum. A mix of automatic and manually selected higher thresholds (see Table S3) were used to correct falsely classified cells. Each cell in the Corrected group is shown by the corrected polar representation and the more strongly filtered Fourier spectrum.

**Figure S5.**
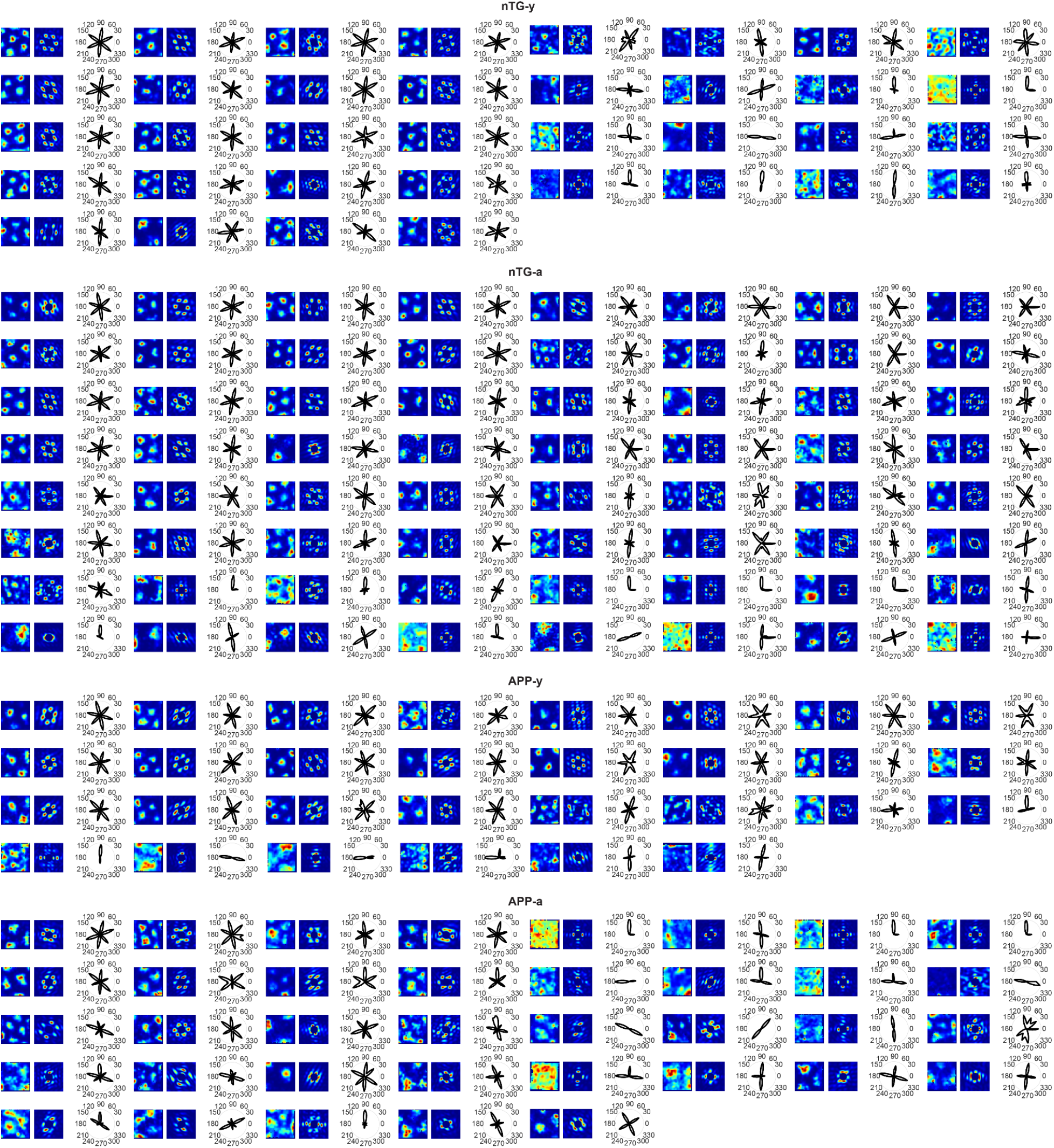
Grid cells, their Fourier components, and polar representations. The spatial rate maps, Fourier components, and polar representations for all grid cells in Figure 3 are shown. The color scale of the rate maps indicates occupancy normalized firing rate. The color scale of the Fourier spectrums indicates Fourier power. The scale of the polar representations indicates Fourier power.

**Figure S6.**
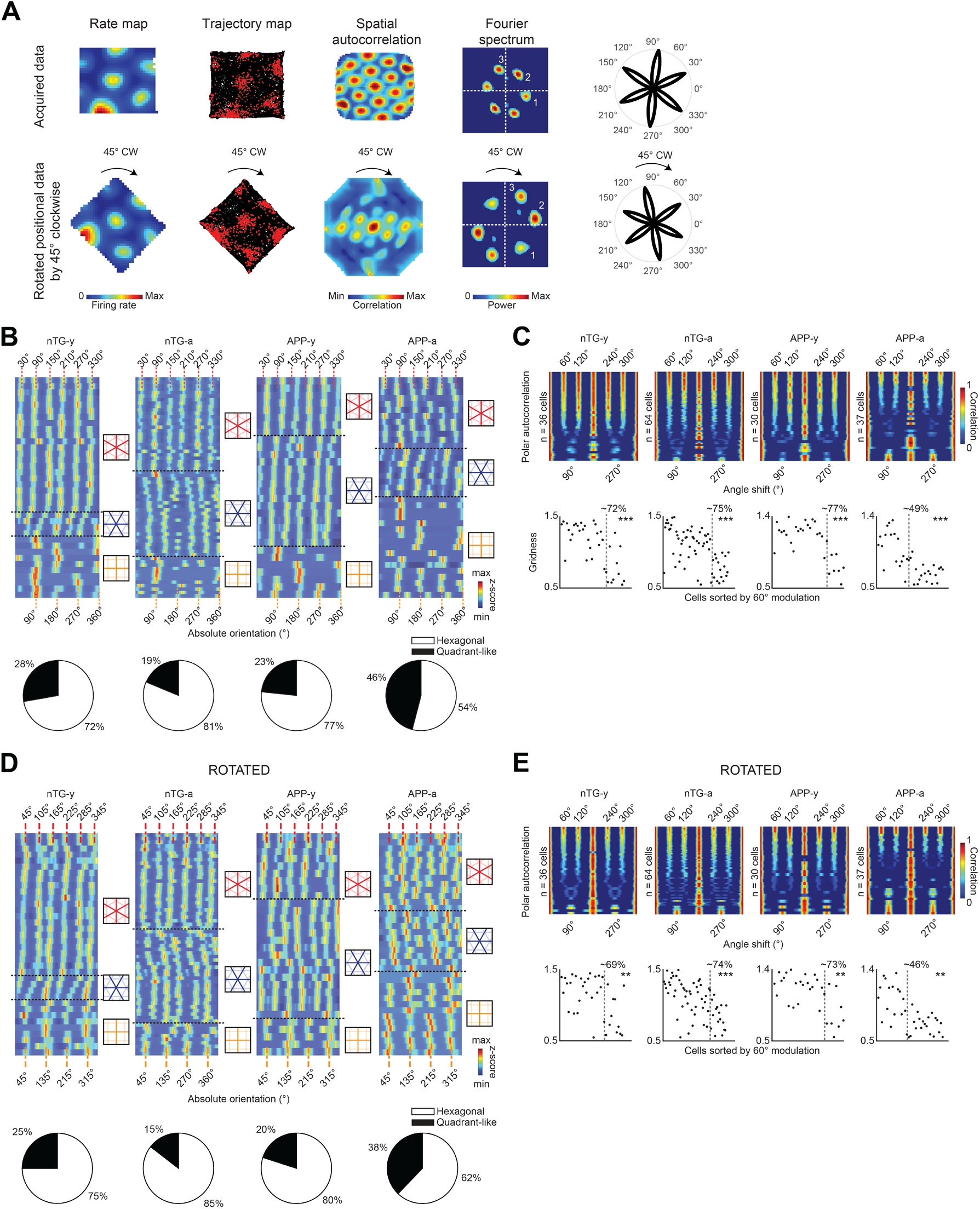
Fourier components reflect rate map content, and not the square shape of images. **(A)** Rotation of positional data by 45° clockwise causes an equal rotation in the Fourier spectrum. **(B)** Top: Absolute orientation of all grid cells relative to the recording environment. Bottom: Percentage distributions of hexagonal or quadrant-like grid cells between groups. **(C)** Polar autocorrelation for grid cells in all groups. Grid cells are sorted by the strength of hexagonal modulation at 60° multiples. Bottom: Gridness scores of grid cells sorted by their degree of hexagonal or quadrant-like modulation along the x-axis. The boundary between the two categories is indicated by the gray dotted lines, and the percentage of strongly hexagonally modulated grid cells are indicated for each group. Wilcoxon rank sum tests. **(D-E)** Same as **B-C**, but for data when positional data was rotated by 45° clockwise. For panels C and E, n.s = non-significant, **p < 0.01, ***p < 0.001.

## Materials and methods

### Mice

J20 APP male mice (B6.Cg-Zbtb20 Tg(PDGFB-APPSwInd) 20Lms/2Mmjax) were obtained from The Jackson Laboratory (MMRRC stock #34836) and bred with female C57/BL6/j mice. Mice were individually housed on a 12-h light/dark cycle and underwent experiments during the light cycle. Housing room conditions of the mice were maintained at 20-22 degrees Celsius and 21-30% humidity. All experimental procedures were performed in accordance with McGill University and Douglas Hospital Research Centre Animal Use and Care Committee (protocol #2015-7725) and in accordance with Canadian Institutes of Health Research guidelines.

J20 mice undergo progressive neuronal loss in layers 2, 3 and 5 of the MEC. By 7.5 months of age, all layers exhibit a total loss of 16.3% of neurons compared to control mice, along with reduced density of presynaptic terminals by 7 months of age (Nagahara et al., 2013). By 6 months of age, region CA1 of the hippocampus also experiences 10%+ of neuronal loss compared to control mice. In parallel, marker-immunoreactivity and electron microscopy confirm the presence of synapse loss in the CA1 as early as 3 months. Besides neuronal and synaptic loss, the complement-dependent pathway and microglia are upregulated in a manner that is dependent on soluble Aβ oligomeric levels in the hippocampus (Hong et al., 2016). On a related note, gliosis (activated astrocytes) and neuroinflammation (activated microglia) are elevated across age in the hippocampus by 6 months of age (Wright et al., 2013). Furthermore, *in vitro* slice electrophysiology experiments reveal that basal synaptic transmission recorded in CA1 and long-term potentiation in the Schaffer collateral–CA1 synapse are impaired by 3 months of age (Saganich et al., 2006). In terms of oscillatory activity, between 4-7 months of age, gamma oscillations are reduced in the parietal cortices which causes network hypersynchrony and are linked to a reduction in the voltage-gated sodium channel subunit Nav1.1 predominantly found in parvalbumin interneurons (Verret et al., 2012). Such hypersynchrony may be linked to spontaneous nonconvulsive seizure activity between 4-7 months, along with numerous inhibitory deficits in the dentate gyrus (Palop et al., 2007). At the behavioral level, J20 mice exhibit numerous spatial navigation impairments in the Morris water maze, the radial arm maze and a food-foraging path integration task in darkness (Cheng et al., 2007; Wright et al., 2013; Ying et al., 2022). To examine the impact of these Aβ-mediated changes on neural coding during early pathology, we restricted our experiments between 3-7 months of age.

Single-unit recording data in the MEC were collected from 68 APP mice and non-transgenic littermates across four experimental groups: young APP mice (3-4.5 months of age), adult APP mice (4.5-7 months of age), young non-transgenic (nTG) mice (3-4.5 months of age), adult nTG mice (4.5-7 months of age). Thirty-one males and 37 females were used. Ten animals fell into multiple age groups. The male/female ratios were 6:5, 16:16, 9:5, and 11:10 for young APP, adult APP, young nTG, and adult nTG mice respectively.

### Surgery

On the day of surgery, mice were anesthetized using isoflurane (0.5% - 3% in O_2_) and administered carprofen (0.01 ml/g) subcutaneously. Three anchor screws were secured to the skull and a ground wire was positioned either above the cerebellum at midline position or the left visual cortex. A ‘versadrive’ containing four independently movable tetrodes (Axona, Inc) was implanted on top of the right MEC at the following stereotaxic coordinates: 3.4 mm lateral to the midline, 0.25-0.40 mm anterior to the transverse sinus. Tetrodes were gold-plated to lower impedances to 150-250 kΩ at 1 kHz prior to surgery. The versadrive was angled at eight degrees in the posterior direction. Following placement, the versadrive was secured in place using Kwik-Sil and dental acrylic. The ground wire was soldered to the implant, and tetrodes were lowered 1.0 mm from the dorsal surface of the MEC. All surgical procedures were performed in accordance with McGill University and Douglas Hospital Research Centre Animal Use and Care Committee (protocol #2015-7725) and in accordance with Canadian Institutes of Health Research guidelines.

### Neural Recordings

Three days post-surgery, mice were placed on water restriction and maintained at 85% of their *ad libidum* weight throughout experiments. Mice were recorded in a 75 x 75 cm box. As mice explored their environments, water droplets were randomly scattered to motivate the subjects to sample the entire arena. Once mice provided good trajectory coverage, tetrodes were turned until theta rhythmic units were observed which indicated that the tetrodes had entered the MEC. Across days, tetrodes were advanced in increments of 25 microns to sample new putative MEC neurons.

To record spikes and local field potentials, versadrives were connected to a multichannel amplifier tethered to a digital Neuralynx (Bozeman, MT) recording system, and data were acquired using Cheetah 5.0 software (Neuralynx, Inc). Signals were amplified and band-pass filtered between 0.6 kHz and 6 kHz. Spike waveform thresholds were adjusted before each recording and ranged between 35-140 μV depending on unit activity. Waveforms that crossed threshold were digitized at 32 kHz and recorded across all four channels of the given tetrode. Local field potentials were recorded across all tetrodes.

### Histology

Animals were anesthetized with Isoflurane and perfused intracardially using saline and 4% paraformaldehyde. Animal heads were left in 4% paraformaldehyde for 24-72 hours following perfusion, before brains were extracted. Brains sanks in a 30% sucrose solution before being frozen and stored in a −80°C freezer. Sagittal brain sections (40μm) were obtained using a cryostat and Nissl-stained with a Cresyl violet solution. In cases where brain slices came off the glass slides during Nissl-staining, slices were instead mounted using a fluorescent DAPI labeling mounting medium.

Tetrode tracks were characterized to be in the superficial or deep layers based on the location of track tips. Only data collected from tetrodes within the MEC were included in analysis.

### Spike sorting

Single-units were isolated ‘offline’ manually using Offline Sorter 2.8.8 (Plexon, Inc). Neurons were separated based on peak amplitudes and principal component measures of spike waveforms. Evaluations of the presence of biologically realistic interspike intervals, temporal autocorrelations, and cross correlations confirmed single-unit isolation. The experimenter was blind to the age and genotype of the subjects and only well-separated clusters were included in analysis.

### Position, direction, velocity estimation, and rate map construction

Positional data was acquired at 30 frames per second at 720 x 480 pixel resolution (4.9 pixels per cm) using a camera purchased from Neuralynx (Bozeman, MT). The camera was elevated at a height to fully capture the entire recording arena. The estimated position of the animal was the centroid of a group of red and green diodes positioned on the recording head stage. Head direction was calculated as the angle between the red and green diodes. Up to five lost samples due to occlusion of tracking LEDs, or reflections in the environment were replaced by a linear interpolation for both position and directional data. Running velocity was calculated using a Kalman filter. Rate maps were constructed by calculating the occupancy-normalized firing rate for 3cm x 3cm bins of positional data. Data were smoothed by a two-dimensional convolution with a pseudo-Gaussian kernel involving a three pixel (9 cm) standard deviation. In most analyses (when specified), rate maps were resized into squares of size 36 x 36 pixels or 26 x 26 pixels.

### Gridness score

We calculated the gridness score using the same procedure described in Brandon et al. 2011 (Brandon et al., 2011). This metric quantifies hexagonal periodicity in the firing rate map, while also accounting for elliptical eccentricity along one of two mirror lines that exist in a hexagonal lattice structure. Distortion along one of the mirror lines was corrected after determining the major and minor axes of the grid based on the six closest fields to the central peak of the rate map autocorrelogram. The entire autocorrelogram was compressed so that the major axis became equal to the minor axis. Large eccentricities where the minor axis was less than half of the major axis were not corrected. From the compressed autocorrelogram, we extracted a ring of the six closest peaks to the center peak. A rotational autocorrelation of this ring was calculated to observe the periodicity in paired pixel correlations across 180 degrees of rotation. The gridness score was computed as the difference between the highest correlation observed at 30, 90, or 150 degrees of rotation and the lowest correlation observed at 60 or 120 degrees of rotation.

### Directionality

The animal’s head direction was collected in bins of 6 degrees and the number of spikes in each bin was divided by the time spent facing that direction. The mean resultant length (MRL) of the polar plot was taken as a metric of directional selectivity.

### Grid cell and head-direction cell selection

Grid cells and head direction cells were determined via a shuffling procedure. Spike trains from each neuron were randomly shifted in time by at least 30 seconds. We then calculated gridness and directionality measures. This process was repeated 50 times for each neuron, and the 99^th^ percentile of the resulting distribution of scores was determined as the significance criteria for both measures. This resulted in a gridness threshold of 0.54 and directionality threshold of 0.21. Cells that passed these thresholds were characterized as grid cells and head direction cells, respectively.

### Spatial stability analysis

Time-partitioned analysis: Rate maps of size 26 x 26 pixels were created for each grid cell. The first 30 minutes of each recording was divided into two 15-minute partitions. Each rate map was divided into wall or center regions, depending on how many pixel layers each had. Next, the pixels corresponding to both regions were extracted for both partitions. A Pearson correlation was computed for wall or center time partitions.

Occupancy-partitioned analysis: A 26 x 26 pixel matrix (an ‘occupancy map’) tracked the number of time frames that the animal spent in each spatial bin throughout the full recording. All time frames where the mouse was stationary (velocity < 5cm/s) were dropped. The values in each bin were then halved, and the resulting numbers represented the number of time frames per bin allocated to each partition. Time frames were then selected throughout the entire recording session in an ‘ascending’ or ‘monotonically increasing’ manner to build both partition maps. Rate maps of size 26 x 26 pixels were created for each partition. The rest of the analysis was the same as the time-partitioned analysis.

### Generation of Fourier spectrums

We followed the procedure used previously in Krupic et al. 2012 (Krupic et al., 2012). We used the fft2 MATLAB function to compute the two-dimensional Fourier transform of the rate map of a given cell. Initial rate maps were unsmoothed at size 36 x 36 pixels and center zero-padded to have size 256 x 256 pixels in order to increase the spatial resolution of the Fourier spectrogram. After running fft2 on the zero-padded rate map, the resulting amplitude spectrum was divided by the mean firing rate of the given cell to allow for comparison between cells that have different firing rates. The amplitude spectrum was element-wise squared to re-visualize the spectrum in the power domain.

We used the fftshift MATLAB function to shift low frequencies to the center and high frequencies to the periphery of the spectrum. Lastly, we created a two-dimensional gaussian loss function (where the bump values are 0) with a width of 1. This gaussian was centered at the strongest value pixel of the spectrum and element-wise multiplied. This procedure was done to erase the center block of energy that resulted from the fft2 function.

### Identifying Fourier components

The Fourier power spectrum was first computed. To reduce effects of noise, the 75^th^ percentile value of power in shuffled data (from the Voronoi procedure) was subtracted and negative values set to zero. To further control for noise, values lower than 25% of the resulting maximum power were also set to zero. These are higher thresholds than what was used by Krupic et al. 2012 (Krupic et al., 2012). However, to ensure that these thresholds do not bias our results, we varied this latter parameter in Table S3.

A zero matrix with the size of the Fourier spectrum was created. Any non-zero value in the corrected Fourier spectrum had its corresponding position in the zero matrix set to 1. Using the regionprops MATLAB function, Fourier components were individually identified as separate “regions”. To account for effects of noise, any regions with an area less than 10 pixels was discarded. Similar to Krupic et al. 2012, cells with more than 4 Fourier components were not included in analysis (Krupic et al., 2012).

To determine a component’s orientation, the distances (along the x- and y-axes) between the centroids of each region and the center of the Fourier spectrum were computed. The “wave vector” corresponding to the x- and y-displacements were calculated as:

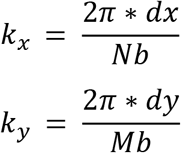

Where dx and dy are the x- and y-displacements, respectively; N and M are the x- and y-axis lengths of the rate map (both 36 in our case); b is the bin size in meters (0.0208 in our data). The orientation of the wave vector was:

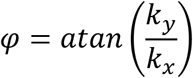

And the orientation of the periodic band (or grid axes) is 90 degrees offset to the wave vector’s orientation.

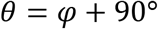

### Fourier polar representations and autocorrelations

To compute Fourier polar representations, we first plotted the orientations of individual Fourier components on a polar plot. The power corresponding to each orientation was that given component’s Fourier power in the Fourier spectrum. The polar plot was then smoothed using a one-dimensional Gaussian kernel with a standard deviation of 13 degrees.

To compute Fourier polar autocorrelations, the smoothed polar plots were circularly shifted 360 degrees. At each degree shift, the Pearson correlation between the original and shifted polar profiles was computed.

### Fourier wavelength identification

To compute a Fourier component’s wavelength, a line was drawn over the Fourier spectrum at the given component’s orientation. This was achieved using the improfile MATLAB function, which draws a line between a specific start (the Fourier spectrum’s center) and end point (a point exceeding the dimensions of the spectrum) on a given image (the spectrum). The distance away from the spectrum’s center along this line which had the maximum Fourier power was taken as the wavelength.

### Fourier orientation offset from reference axes

A grid cell was first determined to be either a 30° or 60° hexagonal grid, or a 90° quadrant-like grid based on where the Fourier orientations faced. The reference axes for a 30° grid faced (30°, 90°, 150°, 210°, 270°, 330°). The reference axes for a 60° grid faced (0°, 60°, 120°, 180°, 240°, 300°). The reference axes for a 90° grid faced (0°, 90°, 180°, 270°). The angular difference between each Fourier component and its closest reference axis was calculated as the offset.

### Theta modulation analyses

To visualize theta modulation, spike-time autocorrelations were computed for each cell with 5ms temporal bins from a lag of −400 to 400ms. The resulting autocorrelations were convolved with a 25ms gaussian and z-score normalized. The average z-scored correlation curve for all cells was also plotted.

To compute intrinsic frequency, the spike-time autocorrelations were zero padded to 2^13 samples and the power spectrum was calculated using the Chronux toolbox function MTSPECTRUMC from Matlab. The intrinsic frequency of a given cell was taken as the frequency with the max power in the 6-12 Hz range. The theta modulation ratio was calculated as the mean power within 6-12 Hz, divided by the mean power between all other frequencies within 2-100 Hz. To be considered a significantly theta-modulated cell, the mean power within 6-12 Hz needed to be four times greater than the mean power within 2-100 Hz (a theta modulation ratio greater than 4).

### Phase precession

The degree of grid cell phase precession was calculated via a ‘pass index’ analysis as described in Climer et al. 2013 (Climer et al., 2013). Briefly, this method quantifies precession by assessing the correlation of a cell’s firing in relation with theta phase as the mouse passes a spatial field. The spatial field locations were estimated using a ‘field index’ method that calculates the degree of occupancy-normalized firing within each bin of positional data. Field contours and centers were then generated by using various parameters involving the field index and firing rate. Lastly, based on the field index signal across a recording session, a mouse’s entry and exit of a spatial field could be estimated. The pass index was therefore determined by normalizing field index signal segments consisting of the passage across a spatial field between −1 and +1, where −1 represents the start of a pass, 0 represents the center, and +1 represents the end. Phase of spiking across field crossing was aligned to this normalized position and a linear-circular correlation was computed between field index and spiking phase (Kempter et al., 2012). A grid cell with a significant correlation at p < 0.05 and a slope per pass between −1440 and −22 was classified as a phase-precessing cell.

### Genotyping

Tail samples were collected at weaning for genotyping, and also prior to brain perfusion. DNA sample were extracted and amplified using REDExtract-N-Amp^™^ Tissue PCR Kit (MilliporeSigma, XNAT-100RXN) and the primer sequence and PCR protocol from The Jackson Laboratory (MMRRC, 34836-JAX). Genotyping results were visualized with a QIAxcel instrument (Qiagen).

### Quantification of neural data

Single-unit data were obtained using Neuralynx (Bozeman, MT) software and isolated ‘offline’ manually using graphical cluster cutting software (Plexon, Inc) individually for each recording session. Custom MATLAB scripts were used to analyze neural data.

### Statistical analysis

Statistical analyses for neural and behavioral data were performed using MATLAB. Bars in bar graphs represent median values, and the error bars represent the 25^th^ and 75th percentile.

Comparisons between two groups involving continuous data used unpaired, two-tailed Wilcoxon rank sum tests. Comparisons between more than two groups involving continuous data used a two-way ANOVA with age and genotype as the factors. A significant interaction effect was followed by post-hoc testing using unpaired, two-tailed Wilcoxon rank sum tests with a Bonferroni-Holm correction in the following four comparisons: nTG-y vs. nTG-a, nTG-y vs APP-y, APP-y vs. APP-a, nTG-a vs. APP-a.

In specific cases where proportions were compared between groups in Figure 2D, a one-tailed t-test for proportions was used to compare 3-component cells, and a two-tailed t-test for proportions was used to compare 2-component cells.

All remaining cases involving proportions in Figures 2F, 3A, 3B, 4C, 4D and 4F used binomial tests. A binomial test determines the probability of an outcome when there are only two possible outcomes. In our case, the test determined if the obtained proportion by a group was equal or unequal (either lesser or larger) compared to expected chance. Expected chance was calculated as the average of the proportions obtained in all four groups. The rationale for calculating expected chance in this manner is assuming that all four animal groups were the same age and genotype, then the average of the four should provide a theoretical level of chance.

All statistical tests used an alpha value of 0.05. Significance was determined as follows: * p < 0.05, ** p < 0.01,*** p < 0.001.

### Data and code availability

All data reported in this paper will be shared by the lead contact upon request. All original code has been deposited at GitHub [insert link] and is publicly available as of the date of publication. DOIs are listed in the key resources table. Any additional information required to reanalyze the data reported in this paper is available from the lead contact upon request.

